# Ketogenic diet dampens excitatory neurotransmission by shrinking synaptic vesicle pools

**DOI:** 10.1101/2025.06.04.657883

**Authors:** Marion I. Stunault, Panyue Deng, Erica M. Periandri, Francisca N. Vitorino, Amelia J. Barfield, Renzelle J. Ponce, Benjamin A. Garcia, Gabor Egervari, Vitaly A. Klyachko, Ghazaleh Ashrafi

**Affiliations:** Department of Cell Biology and Physiology, Washington University School of Medicine, St. Louis, MO, 63132, United States; Department of Genetics, Washington University School of Medicine, St. Louis, MO, 63132, United States; Department of Biochemistry and Molecular Biophysics, Washington University School of Medicine, St. Louis, MO, 63132, United States; Needleman Center for Neurometabolism and Axonal Therapeutics, Washington University School of Medicine, St. Louis, MO, 63132, United States

## Abstract

**HIGHLIGHTS:**

- KD rewires gene expression in the hippocampus, particularly impacting synaptic genes.
- KD profoundly alters epigenetic modifications in the hippocampus.
- KD reduces excitatory synaptic gain and dampens integration of synaptic inputs.
- KD reduces the readily releasable vesicle pool in excitatory synapses.

The ketogenic diet (KD) is a common dietary intervention for treating seizures in intractable childhood epilepsies and has been proposed to improve disease outcome in neurodegenerative disorders. Despite its clinical applications, we know little about how this diet impacts brain circuitry and neuronal function to elicit its protective effects. Here, we examined the impact of the KD on hippocampal function through integrative analysis of gene expression, epigenetics and neurotransmission. We found that KD induces profound transcriptional reprogramming of the hippocampus, including dampened expression of numerous synaptic genes. Through proteomic analysis of histone variants and post-translational modifications, we uncovered significant changes in activating and repressive histone marks in the hippocampus of KD mice. To determine how transcriptional rewiring of the hippocampus under KD impacts neurotransmission, we performed electrophysiological recordings of neurotransmission and synaptic dynamics at excitatory CA3-CA1 synapses. We found that KD diminishes synaptic gain and dampens short-term plasticity at excitatory synapses, resulting in reduced integration of synaptic inputs at the circuit level. Combining electrophysiology and electron microscopy, we determined that effects of KD in excitatory synapses are caused by a reduction in size of the readily releasable pool of synaptic vesicles, as well as the total vesicle pool. Our findings show that the ketogenic diet triggers synaptic remodeling in the hippocampus, driven by broad transcriptional and epigenetic changes that reduce synaptic vesicle pools and short-term plasticity at excitatory synapses ultimately dampening excitatory synaptic gain and integration at the circuit level. These synaptic adaptations may represent a major mechanism underlying the anti-epileptic effects of this diet.

## INTRODUCTION

Ketogenic diets (KDs) are high-fat, low-carbohydrate, and moderate-protein dietary regimens that have been used for nearly a century to treat intractable childhood epilepsies, which account for approximately 30% of all pediatric epilepsy cases^1^. More recently, KDs have been shown to improve disease outcome in various neurological disorders, including Alzheimer’s and Parkinson’s disease^2,3^, schizophrenia^4,5^, and stroke^6,7^. Starvation, and low carbohydrate diets such as KDs, induce a metabolic state known as ketosis^8,9^ in which the liver breaks down lipids and releases ketone bodies (mainly β-hydroxybutyrate (BHB) and acetoacetate) into the bloodstream^10^. In children placed on KD, the gradual increase in circulating ketone bodies coincides with seizure control, which starts within days of the diet, reaching full effects within 1-2 months^11,12^. Despite their clinical potential, the strict dietary restrictions required for KDs pose major challenges for patient adherence, thereby limiting their therapeutic effectiveness. These limitations underscore the need for alternative therapies that harness the neuroprotective mechanisms of KDs. However, there is a significant gap in our understanding of the cellular and molecular processes underlying the anti-epileptic and neuroprotective effects of KDs.

Circulating ketone bodies not only serve as an alternative energy source for various organs, including the brain, but also function as signaling molecules modulating epigenetics through post-translational modifications of histones^13^. The metabolic switch to ketosis exerts both acute and chronic effects on the brain, altering metabolism and neuronal activity over different time scales^14,15^. Indeed, acute neuronal application of ketone bodies is shown to suppress excessive firing by causing membrane hyperpolarization through opening of K_ATP_ channels^16^, and inhibition of glutamate loading into synaptic vesicles (SVs). Altogether, these effects dampen glutamate release during excitatory neurotransmission^17^, contributing to the anti-epileptic properties of KD. The nervous system also needs to adapt to prolonged dietary restrictions which may impact neuronal and circuit activity. However, the long-term effects of KDs on the brain are not fully understood. A handful of studies suggest that KD or administration of ketone bodies induces the expression of brain-derived neurotrophic factor (BDNF)^18^, and alters steady-state levels of brain neurotransmitters, including GABA and glutamate^19,20^. Furthermore, KD profoundly alters gene^21^ and protein^14^ expression across the brain, but the physiological significance of these changes for neurotransmission and circuit function remains largely unknown.

Here, we took a multi-disciplinary approach to uncover the effects of KD on gene expression, epigenetics, and neurotransmission within the hippocampus, an area of the brain that is frequently involved in epileptic seizures and is highly responsive to KDs^22,23^. We found significant downregulation of synaptic gene expression, and major changes in histone modifications in the hippocampus following three weeks of a KD. Electrophysiological analysis of short-term plasticity of hippocampal excitatory synapses revealed reduced synaptic gain and increased short-term synaptic depression in KD mice consistent with a reduction in the readily releasable pool (RRP) of synaptic vesicles in nerve terminals. We conclude that KD triggers profound remodeling of hippocampal synapses, impacting short-term plasticity and neuronal excitability which likely account for the anti-epileptic properties of the diet.

## RESULTS

### A short-term KD alters the expression of synaptic genes in mouse hippocampus

The hippocampus is strongly implicated in intractable epilepsies^24–26^, and hippocampal ketone levels have been shown to rise during ketosis^22^. Therefore we focused on studying the effects of KD on this particular brain region. Metabolic adaptations to nutrient availability often involve changes in gene expression patterns^27^, therefore we investigated transcriptional rewiring of the hippocampus in response to the KD^21^. The effects of KDs on gene expression are often studied after several months of the diet, but we hypothesized that transcriptional rewiring constitutes an early response to dietary intervention, emerging during the initial establishment of ketosis. To determine the onset of ketosis, we monitored blood levels of BHB in 4-week-old C57BL/6NJ male mice placed on a KD (75% fat, 3% carbohydrates, and 8.6% protein) (**Fig. 1A**). While blood BHB level did not change in control mice placed on a chow diet (11.3% fat, 53.4% carbohydrates, and 21% proteins), it increased from 0.4 mM to 1.3 mM within the first week of the KD. Elevated BHB levels were stabilized after three to four weeks of the diet, indicating the establishment of a state of ketosis in KD mice (**Fig. 1B**). Consistent with previous reports^28,29^, the body weight of KD mice remained lower than chow-diet mice over the duration of the diet (**Fig. S1A**).

**Figure 1.**
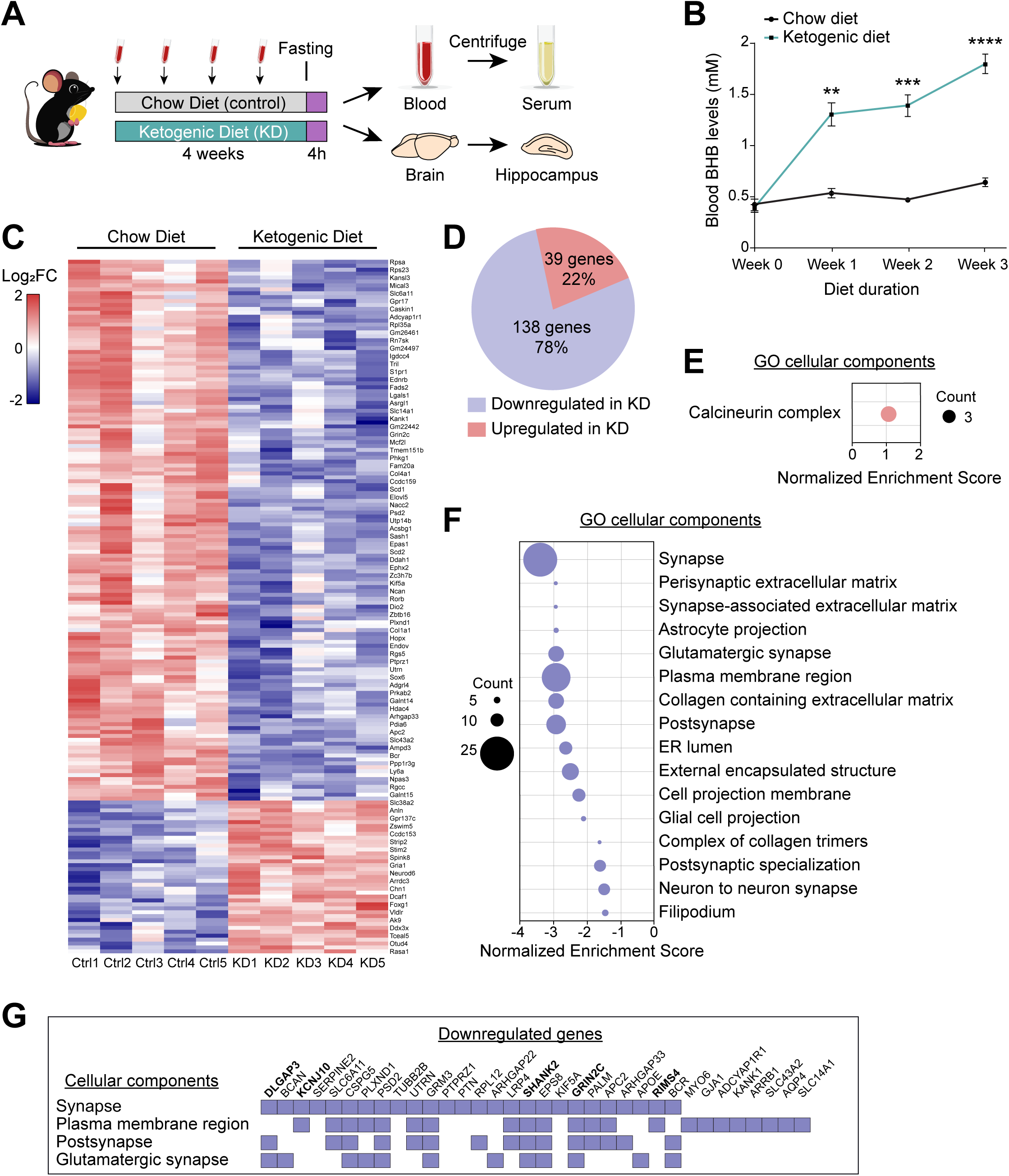
Ketogenic diet induces transcriptional rewiring of mouse hippocampus. **(A)** Schematic of experimental workflow for gene expression analysis. 12-week-old male mice were maintained on either a chow diet or a ketogenic diet (KD) for four weeks. Blood and hippocampus samples were collected for metabolite analysis and RNA sequencing after a four-hour fasting period, respectively. **(B)** Blood β-hydroxybutyrate (BHB) levels over the course of the diets. Not significant: p >0.05, **, 0.01≥ p >0.001; ***, 0.001≥ p >0.0001; ****, p ≤0.0001. Data are represented as mean ± SEM. **(C)** Heatmap showing differential expression of hippocampal genes in mice on chow diet or the KD. Log2 fold change (Log_2_FC) indicates magnitude of gene expression changes in the KD group. Blue, downregulated; red, upregulated. **(D)** Pie chart showing the proportion of genes that were downregulated (78%, 138 genes) and upregulated (22%, 39 genes) in KD mice compared to chow-diet controls. **(E)** Gene Ontology (GO) analysis identifying cellular components significantly associated with the upregulated genes. Circle size corresponds to gene counts. Normalized enrichment score corresponds to -log10(adjusted-p-value). **(F)** GO cellular component analysis of downregulated genes showing enrichment of synaptic structures, including perisynaptic extracellular matrix and post-synapse. Circle size corresponds to gene count. Normalized enrichment score corresponds to -log10(adjusted p-value). **(G)** Table showing cellular component analysis of downregulated genes involved in synapse, plasma membrane region, post-synapse, and glutamatergic synapse. KD, ketogenic diet. N = 5 animal per diet.

We then performed RNA sequencing on hippocampi of adult male mice placed on KD or chow diet for four weeks (**Fig. 1A**). Due to individual variations in food intake, mice were fasted for four hours prior to tissue extraction for RNA sequencing. Postmortem analysis of serum metabolites revealed similar glucose and fatty acid levels in both groups while BHB and triglyceride levels were significantly higher in KD mice (**Fig. S1B-S1E**). Cluster analysis of differentially expressed genes showed that KD mice formed a cohesive cluster, distinct from chow-fed controls, indicating robust modulation of transcriptional activity by KD (**Fig. 1C**). We found 177 genes (false discovery rate<0.05) that were differentially expressed between the two groups, of which 138 genes (78% of all differentially expressed genes) were downregulated and 39 (22%) were upregulated in KD mice (**Fig. 1D**). Pathway analysis of upregulated genes revealed significant enrichment of Gene Ontology (GO) cellular components related to the calcineurin complex (**Fig. 1E**). The genes downregulated in KD mice were predominantly associated with neuronal synapses, particularly glutamatergic (excitatory) synapses, as well as perisynaptic extracellular matrix and astrocyte projections (**Fig. 1F**). Particularly, these genes encode key synaptic components, including presynaptic proteins like RIMS4, which regulates synaptic vesicle exocytosis, and postsynaptic proteins such as DLGAP3 and SHANK2, involved in synaptic scaffolding and signaling (**Fig. 1G**). The downregulated genes also include *Kcnj10* (*Kir4.1*) and *Grin2c* which encode a potassium channel and a component of the NMDA receptor, respectively. Overall, our gene expression analysis indicates that the KD drives synaptic remodeling, with important implications for neurotransmission.

Since KD is often used for treatment of intractable epilepsies in children, we also examined hippocampal gene expression in young (four-week-old) mice placed on the KD. As with adult mice, young KD mice had consistently lower body weight compared to chow-fed mice (**Fig. S2A**). Analysis of serum metabolites after three weeks on the diet confirmed higher BHB and lower glucose levels in young KD mice as compared to chow-diet mice (**Fig. S2B-S2E**). Although we found fewer differentially regulated genes in young mice (63 vs. 177) compared to adult mice (**Fig. S2F and S2G**), most of these genes (60%) were downregulated, as was observed in adult mice (**Fig. 1D**). Furthermore, downregulated genes in young KD mice included those involved in synaptic plasticity and neuronal excitability, namely, *Npas4*, *Kcnj2*, and *Nptx2* (**Fig. S2H**). Therefore, we conclude from our gene expression analysis that following three weeks of the KD, the hippocampus undergoes substantial transcriptional rewiring of synaptic gene expression in both young and adult mice.

### The KD reshapes the epigenetic landscape of mouse hippocampus

Metabolic regulation of the epigenome has emerged as a critical mechanism for fine tuning gene expression programs in the brain^30^, particularly in response to external stimuli. In KD animals, ketone bodies are not only metabolized in the brain but also serve as substrates or co-factors for epigenetic modifiers^31^ and thereby shaping histone post-translational modifications (hPTMs), gene expression, and behavior in the brain^32^. To assess how KD impacts the hippocampal epigenome, we performed liquid chromatography coupled with tandem mass spectrometry (LC-MS/MS) to analyze the relative abundance of hippocampal histone post-translational modifications (hPTMs). Four-week-old male mice were placed on chow or KD diet for three weeks, and their hippocampi were collected for histone extraction (**Fig. 2A**). Agglomerative hierarchical clustering of histone peptides detected by LC-MS/MS revealed a clear separation of hPTM enrichment profiles in hippocampi of chow-fed versus KD mice (**Fig. 2B**). The average relative abundance of 22 histone peptides (7 down, 15 up) were significantly altered in KD mice as compared to chow-diet mice (**Fig. 2C**). These included hPTMs (both acetylation and methylation) of canonical histone H3, which have been tightly linked to gene regulation^33^, as well as overall levels and PTMs of several histone variants (**Fig. 2D**). Specifically, we observed increased acetylation of the H2A.Z variant (**Fig. 2D**), which has been linked to hippocampal gene expression and long-term memory formation, particularly by impacting the expression of synaptic genes^34,35^.

**Figure 2.**
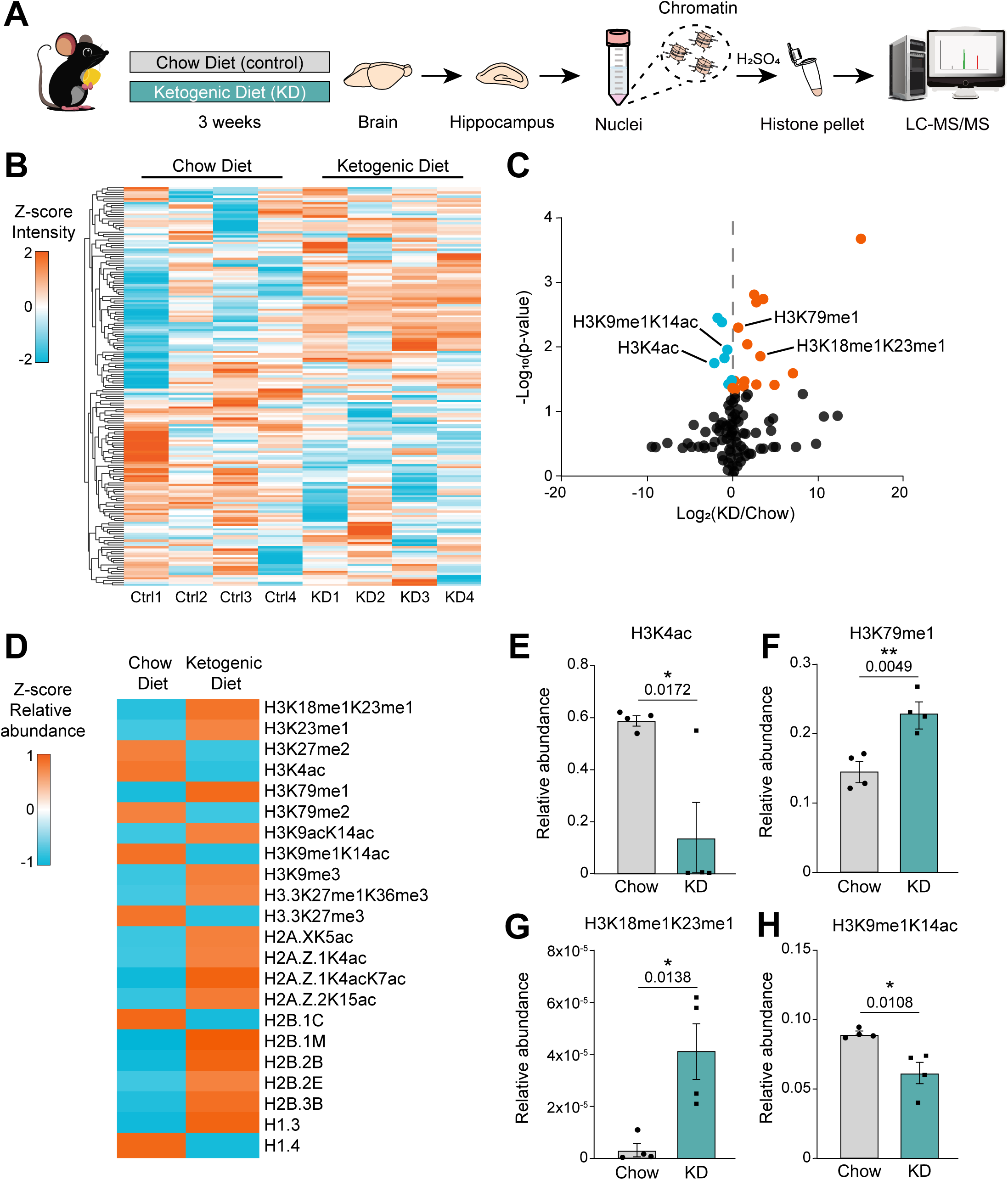
Ketogenic diet alters the epigenetic landscape of mouse hippocampus. **(A)** Schematic of experimental workflow for analysis of epigenetic marks. Four-week-old male mice were maintained on either a chow diet or a ketogenic diet (KD) for three weeks. Hippocampi were harvested and histones were isolated for proteomic analysis by liquid chromatography and mass spectrometry. **(B)** Intensity heatmap of histone peptides in the hippocampus of KD or chow-diet mice, generated with Mass Dynamics^78^ using agglomerative hierarchical clustering. **(C)** Volcano plot of histone peptides in hippocampi of KD and chow-diet mice. Dots represent individual histone peptides (gray: not significant, teal: significantly downregulated, orange: significantly upregulated in KD vs. chow-diet mice). Not significant, p >0.05. **(D)** Heatmap displaying average row Z-scores of relative abundance for histone peptides significantly altered in KD hippocampi as compared to chow diet. **(E-H)** Average relative abundance of a subset of significantly changed histone post-translational modifications (hPTMs). Not significant: p >0.05. Data are represented as mean ± SEM. **(E)** Average relative abundance of H3K4ac (two-tailed unpaired t test with Welch’s correction t_3.115_=3.263, p=0.0446). **(F)** Average relative abundance of H3K9me1K14ac (two-tailed unpaired t test with Welch’s correction t_3.273_=3.645, p=0.0308). **(G)** Average relative abundance of H3K79me1 (two-tailed unpaired t test of equal variance t_6_=4.336, p=0.0049). **(H)** Average relative abundance of H3K18me1K23me1 (two-tailed unpaired t test with Welch’s correction t_3.333_=3.443, p=0.0350). N = 4 animals per diet. Ac, acetyl; hPTMs, histone post-translational modifications; KD, ketogenic diet; me1, monomethyl; me2, dimethyl; me3, trimethyl.

Importantly, we found that KD altered the levels of both activating and repressive histone marks in the hippocampus. For example, the average relative abundance of H3K4ac was significantly reduced in KD mice (**Fig. 2E**). H3K4ac is an hPTM enriched at active gene promoters^36^, and loss of this mark could contribute to the predominant downregulation of gene expression observed in KD (**Fig. 1**). Further, we found significantly increased levels of H3K79me1 and H3K18me1K23me1 peptides in the hippocampus of KD-fed mice (**Fig. 2F** and **2G)**. Mono-methylation of H3K79^37^, H3K18me^38^ and H3K23me^39^ have all been linked to decreased gene expression in various contexts. On the other hand, we found that the average relative abundance of H3K9meK14ac was reduced in the hippocampus of KD-fed mice (**Fig. 2H**). H3K9meK14ac has been implicated in the silencing of active genomic regions by recruiting the repressive K9 methyltransferase SETDB1^40^. The loss of this repressive histone mark could play a role in the increased expression of a small subset of genes observed in our RNA-seq (**Fig. 1C-E**).

In summary, we found that the KD had a marked effect on the hippocampal epigenome, resulting in altered enrichment of both activating and repressive hPTMs as well as histone variants. These epigenetic changes (**Fig. 2**) are consistent with the global transcriptional effect (**Fig. 1**) of KD in the hippocampus.

### KD dampens synaptic gain and reduces the readily releasable vesicle pool in excitatory hippocampal synapses

The effects of KD on global transcription, particularly components of pre- and post-synapse, suggest a potential impact of KD on hippocampal synaptic transmission. Therefore, we performed electrophysiological recordings in acute slices prepared from chow-diet or KD mice to investigate excitatory neurotransmission in hippocampal CA3-CA1 synapses formed by the Schaffer collaterals onto CA1 pyramidal neurons (**Fig. 3A**). We assessed both baseline synaptic transmission evoked by low-frequency (1Hz) stimulation, and its dynamics during high-frequency (40Hz) trains (**Fig. 3B**). We particularly focused on rapid activity-dependent dynamics of neurotransmission, also known as short-term plasticity (STP), which is widely believed to play an essential role in information processing^41,42^. Whole-cell patch-clamp recordings from CA1 pyramidal neurons were performed at near-physiological temperatures (33-34°C) and in the presence of Gabazine to block inhibitory responses. Excitatory postsynaptic currents (EPSCs) evoked during stimulus trains were normalized to an average of five low-frequency controls preceding each train, thus representing relative changes in synaptic strength, i.e. synaptic gain. We observed that while baseline synaptic transmission was largely unaffected in KD mice (**Fig S3A**), synaptic gain during 40Hz trains was significantly reduced (**Fig. 3B-D**) (Chow: 2.52 ± 0.15; KD: 1.95 ± 0.16).

**Figure 3.**
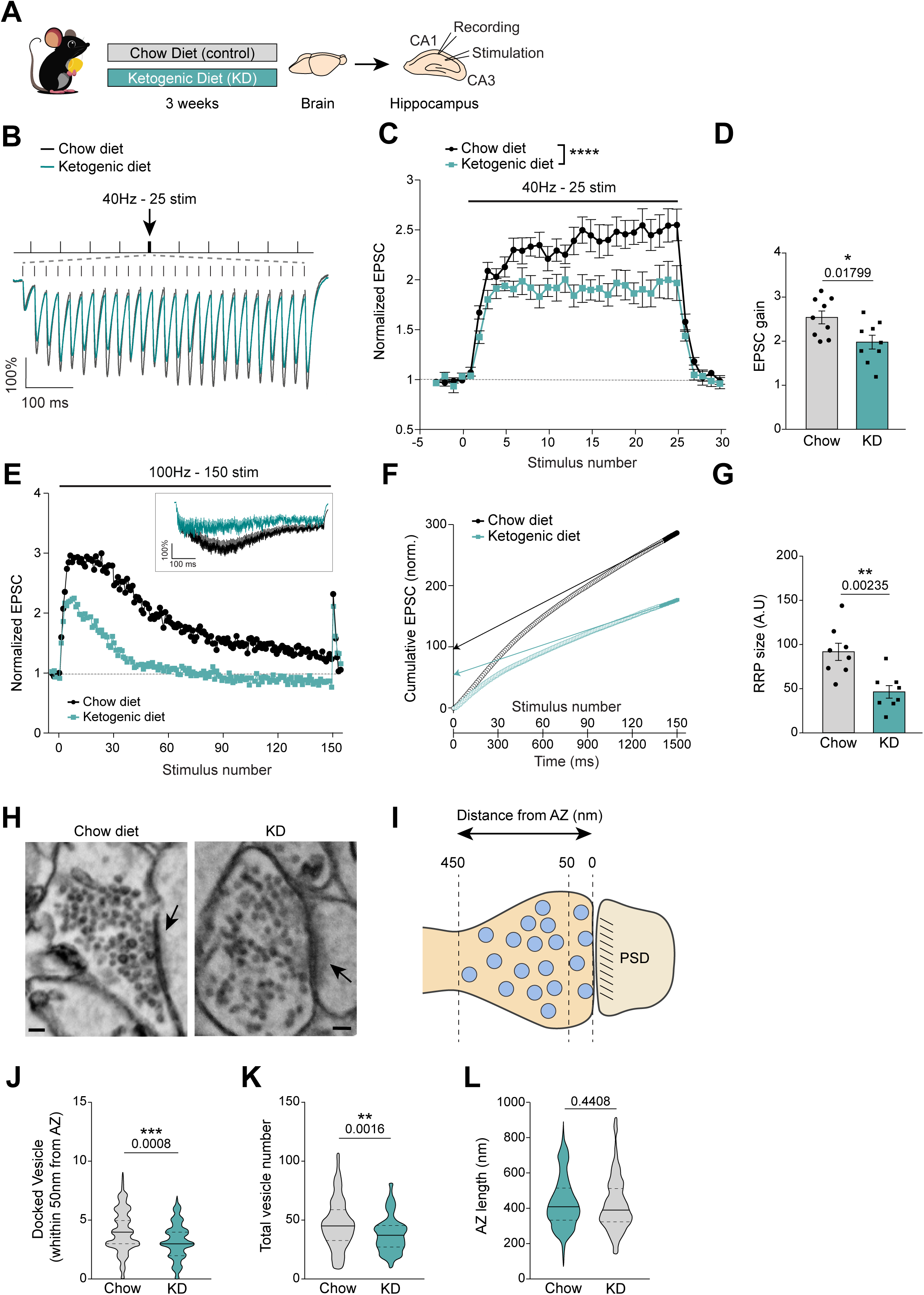
Ketogenic diet dampens excitatory neurotransmission by shrinking the readily releasable pool of synaptic vesicles. **(A)** Schematic of the experimental workflow for acute slice recordings. Four-week-old male mice were maintained on either a chow diet or ketogenic diet (KD) for three weeks. EPSCs were recorded from CA1 pyramidal neurons by stimulating Schaffer collaterals in the presence of Gabazine to block inhibition. **(B, C)** Sample EPSC traces **(B)** and normalized EPSC amplitudes **(C)** recorded in CA1 pyramidal cells in response to Schaffer collateral stimulation with a train of 25 stimuli at 40 Hz (black bars), with Gabazine to block inhibitory responses. EPSC traces were normalized to their own baseline amplitude measured at 0.2 Hz of the same train for better visual comparison. The first EPSCs during the burst in each condition were scaled to their baseline values, representing 100% shown as the vertical bar. Stimulation artifacts were removed for clarity. ****, p ≤0.0001. **(D)** Quantification of EPSC synaptic gain during train stimulation. N = 9 cells/3 animals per diet. Not significant: p >0.05. Data are presented as mean ± SEM. **(E)** EPSC dynamics and representative EPSC traces (Insert) recorded in CA1 pyramidal cells in response to Schaffer collateral stimulation with a train of 150 stimuli at 100 Hz to deplete the readily-releasable vesicle pool (RRP). **(F, G)** Cumulative normalized EPSCs during 150 stimuli at 100 Hz to estimate RRP size. The last 10 cumulative EPSC points were fitted linearly and extrapolated to the time = 0 to estimate RRP size (arrows) **(F)**, and estimated RRP size **(G)**. N = 8 cells/3 animals per diet. Not significant: p >0.05. Data are represented as mean ± SEM. **(H)** Representative electron micrographs of synapses located in the stratum radiatum of the CA1 region of the hippocampus from chow diet-fed mice and KD mice. Black arrows indicate the post-synaptic density (PSD). Scale bars = 100nm. **(I)** Schematic showing vesicle distribution within a synapse. Vesicles located within 50nm of the active zone (AZ) are considered as docked. PSD: Post-synaptic density. **(J, K, L)** Quantification of the docked vesicles (within 50nm of the AZ) **(J)**, total synaptic vesicle number **(K)** and AZ length **(L)** of synapses in the CA1 region of the hippocampus from chow diet-fed mice (control) and KD mice. N = 93 synapses/3 animals (chow); 105 synapses/3 animals (KD). Not significant: p >0.05. Data are represented as mean ± SEM.

These rapid changes in synaptic strength during high-frequency activity, which are known as short-term plasticity (STP), are determined by the interplay of several processes that act to counterbalance each other, namely facilitation working to increase synaptic gain and short-term depression working to decrease it. Facilitation can be assessed using a paired-pulse protocol, however we found no significant changes in the paired-pulse facilitation in KD mice (**Fig S3C-E)**. The attenuation of synaptic gain that we observed in KD mice could thus arise from increased short-term synaptic depression, a fundamental mechanism of STP which is determined predominantly by the availability of synaptic vesicles for release. Therefore, we hypothesized that a decrease in the synaptic vesicle pool known as readily releasable pool (RRP), comprising vesicles docked at the synaptic plasma membrane and immediately available for release, could be responsible for the diminished STP in KD mice. A functional estimate for the size of the RRP can be obtained using analysis of cumulative release during steady-state depression evoked by prolonged high-frequency trains when the release is predominately limited by re-supply of vesicles^43^. In this case, the linear back-extrapolation of the steady-state EPSC amplitude at the end of a high-frequency train has been shown to be proportional to the RRP size^43^. Thus, to probe the size of the RRP in KD mice, we applied a train of 150 stimuli at 100Hz to evoke steady-state depression (**Fig. 3E insert**), which was indeed notably increased in KD compared to chow-diet mice (**Fig. 3E**). However, baseline synaptic transmission was largely unaffected in KD mice (**Fig S3B**). Analysis of cumulative release (**Fig. 3F**) using this approach reveals a significantly smaller functional RRP at CA3-CA1 excitatory synapses in KD mice (**Fig. 3G**), consistent with our hypothesis. Structurally, the RRP consists of synaptic vesicles docked at the plasma membrane of the active zone (AZ)^44^. To validate KD-induced reduction in RRP size, we used electron microscopy to examine synaptic vesicle populations in hippocampal slices from both chow-fed and KD mice (**Fig. 3H**). Docked vesicles are typically defined as those localized within 50 nm of AZ plasma membrane^45^, which along with vesicles positioned more distally constitute the total synaptic vesicle pool in nerve terminals (**Fig. 3I**). Quantification of vesicle populations in CA3-CA1 synapses within the stratum radiatum revealed significant reduction in the number of docked vesicles (Chow: 4.03 ± 0.19; KD: 3.24 ± 0.14) as well as total vesicle pool (Chow: 46.51 ± 2.14; KD: 38.37 ± 1.47) in KD mice (**Fig. 3J, and 3K**). In contrast, the size of the AZ remained unchanged in KD hippocampal synapses as compared to chow-fed mice (**Fig. 3L**). Our functional and morphological observations together suggest that KD causes a reduction in RRP size, which leads to diminished synaptic gain and STP in excitatory hippocampal synapses.

### KD attenuates integration of synaptic inputs in the hippocampal circuit

In addition to directly targeting CA1 pyramidal neurons, Schaffer collateral inputs from the CA3 project onto CA1 inhibitory interneurons, which in turn provide feed-forward inhibition onto CA1 pyramidal cells (**Fig 4A**). Activation of CA1 pyramidal neurons also provides excitatory drive to the local interneurons that produce feed-back inhibition (**Fig 4A**). KD-induced dampening of excitatory gain in CA1 synapses and the resulting changes in excitability of CA1 neurons may therefore alter both the excitatory and inhibitory components of the circuit and change the excitation/inhibition circuit balance in a complex way. To examine the combined circuit-level effects of KD on CA1 pyramidal cell excitability, we examined summation of excitatory postsynaptic potentials (EPSPs) in CA1 pyramidal cells evoked by trains of Schaffer Collateral stimulation. This temporal summation is a critical determinant of neuronal excitability that integrates all of the direct excitatory and indirect inhibitory inputs in the local circuit, including both feed-forward and feed-back inhibition evoked by Schaffer collateral stimulation. A stimulus train of five stimuli at 40 Hz was used to measure EPSP summation in the absence of any glutamate or GABA receptor blockers to keep both excitatory and inhibitory pathways intact. Stimulation electrodes were placed at least 300 μm away from the recorded cells to avoid directly stimulating inhibitory inputs. We observed that KD significantly dampened the EPSP summation (**Fig. 4B** and **4D**). This was not due to the differences in the stimulation intensity, because the first EPSP amplitude was comparable in both conditions (**Fig. 4C**). We further examined EPSP summation in response to 20 and 100 Hz stimulation. Interestingly, KD had no measurable effect on EPSP summation in the low or high frequency inputs (**Fig. 4E**). These results suggest that the net circuit effect of reduced excitatory synaptic gain in the CA3-CA1 circuit is an overall reduction in excitatory-inhibitory balance in a specific frequency range.

**Figure 4.**
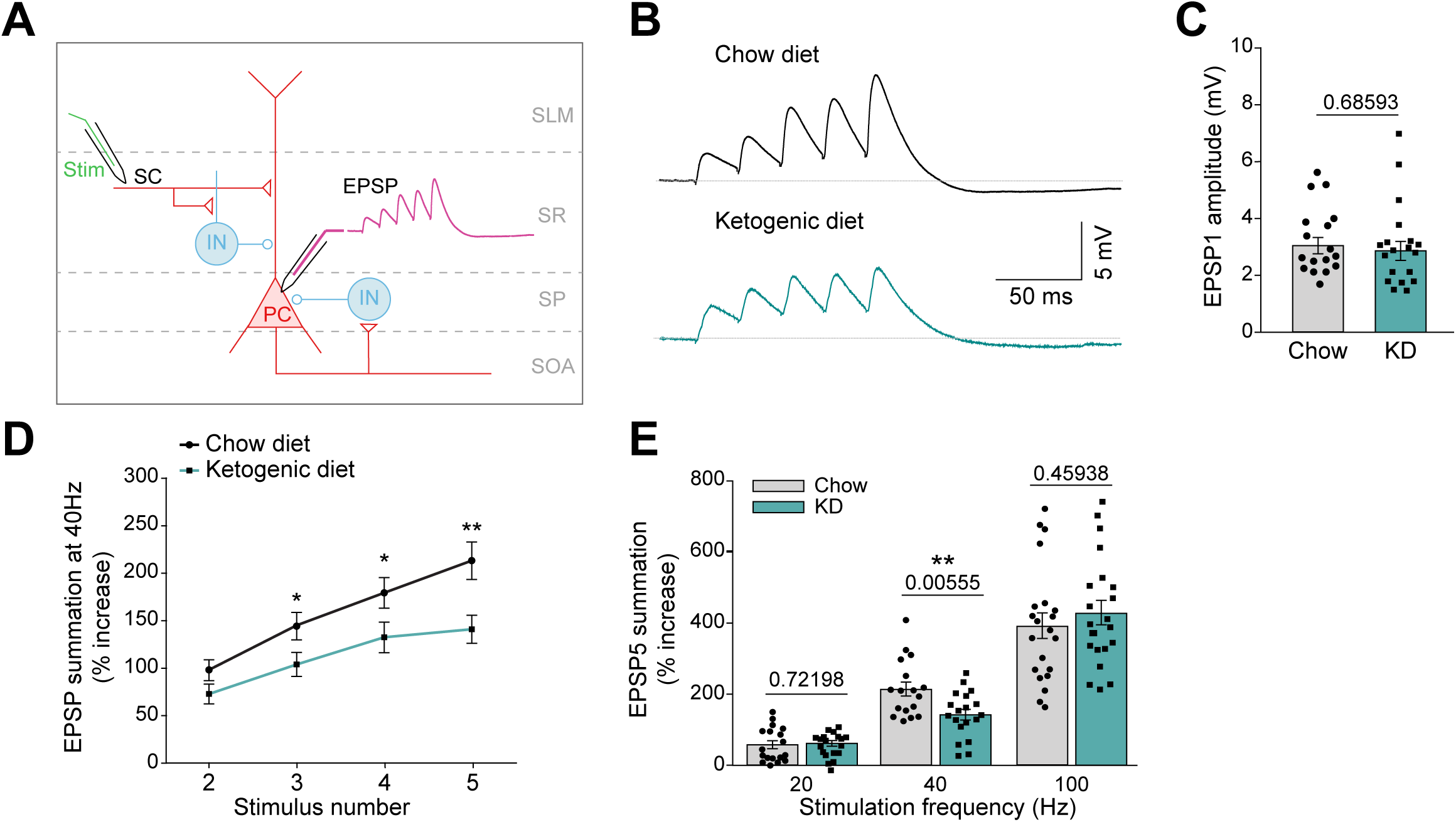
Ketogenic diet reduces EPSP temporal summation in CA1 pyramidal cells. **(A)** Schematic of the CA1 circuit and summation of excitatory post-synaptic potentials (EPSP) in CA1 pyramidal cells. For clarity, the temporoammonic pathway (irrelevant to current study) was omitted and two types of interneurons are shown with axons terminating onto proximate dendrite and soma for feed forward and feedback circuits, respectively. Excitatory (red) and inhibitory (blue) pathways. IN, Interneuron; PC, Pyramidal Cell; SC, Schaffer Collateral; SLM, Stratum Lacunosum-Molecular; SOA, Stratum Oriens-Alveus; SP, Stratum Pyramidal; SR, Stratum Radiatum; Stim, Stimulation. **(B, C)** Sample EPSP traces **(B)**, and quantification of 1st EPSP amplitude **(C)** recorded from CA1 pyramidal cells in response to a train of 5 stimuli at 20, 40 or 100 Hz through an electrode located in stratum radiatum of CA1. N = 17 cells/3 animals (chow); 19 cells/3 animals (KD). **(D)** EPSP summation of 2 to 5 stimuli at 40 Hz. N= 17 cells/3 animals (chow); 19 cells/3 animals (KD). **(E)** EPSP summation of stimulus #5 for stimulation frequencies of 20 (N = 18 cells/3 animals [chow]; 19 cells/3 animals [KD]), 40 Hz (N = 17 cells/3 animals [chow]; 19 cells/3 animals [KD]), and 100 Hz (N = 21 cells/3 animals (chow); 21 cells/3 animals [KD]). *, 0.05≥ p >0.01; **, 0.01≥ p >0.001. Not significant: p >0.05. Data are represented as mean ± SEM.

## DISCUSSION

Therapeutic KD is a highly restrictive dietary regimen with protective effects against epilepsy, psychiatric diseases, and neurodegeneration. Understanding the precise mechanisms underlying these benefits has been challenging due to the complex and pleiotropic effects. The increased reliance of the brain on ketones as an energy source triggers extensive adaptations, both in the short term as ketosis is established, and in the long term as it is maintained. We uncovered that the KD profoundly alters synaptic gene expression in the hippocampus suggesting that synapses are particularly sensitive to the diet. While bulk RNA sequencing can obscure cell type-specific changes in gene expression, the prominence of synaptic signatures in our dataset underscores the specific impact of KD on neuronal transcriptional programs. Synapses may be preferentially remodeled under KD because of their inherently high energy demands and their need to adapt to the metabolic challenges induced by the diet. Future studies will investigate how alterations in the expression of each of these synaptic genes impacts hippocampal functions. Many of the differentially expressed genes in KD hippocampi encode postsynaptic components. Consistent with previous reports^46^, we found that baseline EPSPs were not affected in KD mice (**Fig. S3A, B**). Changes in expression of postsynaptic proteins, such as ion channels and scaffolding proteins may indicate homeostatic adaptations to maintain neuronal network stability under changing metabolic conditions. Further studies are needed to determine how changes in postsynaptic gene expression under KD impact dendritic function, such as signal integration and plasticity. It is also important to investigate cell type-specific transcriptional responses of neurons and glial cells to KD, as metabolic pathways such as glycolysis and OXPHOS are differentially impacted across distinct cell types^14^.

We discovered that KD significantly alters epigenetic modifications in the hippocampus. Ketone bodies such as BHB alter chromatin both by direct deposition on specific histone lysine residues^47^, as well as by regulating the enrichment of canonical histone marks including H3K9 acetylation^48^. Ketone bodies also function as endogenous inhibitors of class I histone deacetylases (HDACs)^49,50^ activating expression of some genes while repressing others^51^. Although we did not observe global elevation of histone acetylation in KD mice, acetylation of the histone variant H2A.Z was increased **(Fig. 2D)**. This finding is particularly intriguing as deposition of H2A.Z on chromatin has been shown to preferentially impact the expression of synaptic genes, and its acetylation is implicated in hippocampal memory formation^35^. While KD profoundly alters histone modifications within the hippocampus, most of these hPTMs have been studied in non-neuronal tissues and their specific role in the mammalian brain remains poorly understood. Therefore, future experiments are needed to link specific histone modifications to gene expression programs by investigating how KD affects the locus-specific enrichment of histone marks across the genome.

How do broad changes in synaptic gene expression ultimately shape hippocampal function? To approach this question, we examined excitatory neurotransmission in hippocampal CA3-CA1 synapses and found a decrease in synaptic gain during high-frequency stimulation along with a shift in STP towards increased short-term depression in KD animals. STP arises from the complex interplay among multiple, interdependent synaptic mechanisms, including several components of synaptic facilitation that work to increase synaptic strength due to accumulation of residual calcium in presynaptic terminals, and opposing short-term depression that is driven primarily by the depletion of vesicles available for release^52^. Our results suggest that reduction of synaptic gain and the shift in STP is driven primarily by smaller RRP size in KD mice, rather than facilitation as indicated by the lack of change in paired pulse ratio.

In the hippocampus, bursts of high-frequency firing generated by place cells are believed to carry spatial information about the environment^53,54^. High-frequency firing of hippocampal excitatory neurons is also believed to represent a response to various non-spatial sensory inputs and to combine sensory information and spatial contexts to process context-specific or episodic information^55^. STP is generally believed to play a central role in these computations^41,56^ and we and others previously showed that STP modulates information processing and transfer in a range of input rates, particularly in the naturally occurring spike frequencies at hippocampal synapses. Interestingly, abnormally elevated STP has been implicated in neurodevelopmental disorders, such as Fragile X syndrome and autism^57,58^, which have a high comorbidity with epileptic seizures. The observed shift in STP towards reduced synaptic gain during bursts of activity may thus have an anti-epileptic effect in part by reducing glutamate release during high-frequency activity and dampening the responsiveness of CA1 neurons, thereby altering the circuit balance between excitation and inhibition. Indeed, we observed that temporal summation in CA1 neurons, which combines both excitatory and inhibitory synaptic inputs, is dampened in KD mice, particularly in a frequency range relevant to the naturally occurring firing of place cells^53,54^. This form of integration is a critical determinant of neuronal excitability, and its dampening in KD mice may thus represent a major mechanism contributing to the anti-epileptic effects of the KD. Previous studies have attribute the anti-epileptic properties of the KD to a shift toward a more inhibitory state, driven by an increased GABA/glutamate ratio in the brain^19^. Our results suggest that presynaptic mechanisms of STP, such as reduction in the RRP size, may also contribute to altered excitation-inhibition balance under KD by increasing short-term depression in excitatory synapses and reducing excitatory synaptic gain. KD has also been suggested to affect long-term potentiation (LTP), a form of synaptic plasticity marked by sustained increase in synaptic strength, but extensive studies of LTP changes in KD mice yielded conflicting outcomes^46,59,60^. In contrast, our study demonstrates that KD has robust effects on STP by reducing the size of synaptic vesicle pools, highlighting this rapid form of plasticity as a key target of the diet.

Our functional and morphological studies of hippocampal presynaptic terminals revealed that readily releasable and total vesicle pools are diminished in KD mice. However, the underlying mechanisms that drive this reduction in vesicle pool size remain unknown. We hypothesize that such synaptic remodeling results directly from changes in expression of synaptic genes, but further work is needed to elucidate the underlying molecular pathways. KD may impact biogenesis, axonal trafficking, or presynaptic docking of synaptic vesicles, therefore it is particularly important to investigate the composition of SNARES and their accessory proteins at the active zone in KD mice. Furthermore, it is also critical to determine whether KD alters vesicle pool size and synaptic strength in other types of synapses such as inhibitory ones, or different brain regions such as dentate gyrus or cortex which are also involved in epileptic seizures. Given the enrichment of excitatory synapse markers in our gene expression dataset, we predict that KD differentially modulates excitatory synapses to re-normalize the excitatory– inhibitory balance in disease states. Future studies are needed to investigate how KD modulates synaptic plasticity and presynaptic architecture across diverse mouse models of epilepsy.

## AKNOWLEDGMENT

We thank the Genome Technology Access Center at the McDonnell Genome Institute at Washington University School of Medicine for help with genomic analysis; J. Sponagel for technical assistance. Metabolites analyses were performed by the Washington University Nutrition Obesity Research Center (NORC), supported by the NIH grant P30 DK056341. We acknowledge the Washington University Center for Cellular Imaging (WUCCI) which is supported by Washington University School of Medicine, The Children’s Discovery Institute of University and St. Louis Children’s Hospital (CDI-CORE-2015-505 and CDI-CORE-2019-813) and the Foundation for Barnes-Jewish Hospital (3770 and 4642). This project was supported by Diabetes Research Center Pilot & Feasibility Award (G.A.) funded by the NIDDK Grant No. P30 DK020579, NIGMS R35GM147222 (G.A.), the Whitehall Foundation (G.A.), the Chan-Zuckerberg Initiative Early Acceleration Award (G.A.), R00AA028577 (G.E.), NARSAD Young Investigator Grant YIG31527 (G.E.), 7R01AI118891 (B.A.G.), R01HD106051 (B.A.G.), and NINDS R35 NS111596 (to V.A.K.).

## Author contributions

M.I.S. and P.D., conceptualization, investigation, visualization, formal analysis, and writing (original draft, review, and editing); E.M.P., F.N.V., A.J.B., and R.J.P., investigation, visualization, formal analysis; B.A.G., G.E., V.A.K, and G.A., conceptualization, funding acquisition, supervision and writing (original draft, review, and editing).

## STAR↔METHODS

| **REAGENT or RESOURCE** | **SOURCE** | **IDENTIFIER** |
|---|---|---|
| **Animals** |  |  |
| C57BL/6NJ | Jackson laboratory | Strain #005304 |
| **Chemicals, Peptides, and Recombinant Proteins** |  |  |
| 4-(2-Hydroxyethyl)piperazine-1-ethanesulfonic acid, N-(2-Hydroxyethyl)piperazine-N′-(2-ethanesulfonic acid) (HEPES) | MilliporeSigma | Cat. No. H3375 |
| Adenosine 5'-triphosphate disodium (Na2-ATP) | MilliporeSigma | Cat. No. A1852 |
| CA-630 | Sigma-Aldrich | Cat. No. I8896 |
| D-(-)-2-Amino-5-phosphonopentanoic acid (APV) | Tocris Bioscience | Cat. No. 0106 |
| Guanosine 5′-triphosphate sodium (Na-GTP) | MilliporeSigma | Cat. No. G8877 |
| Sequencing-grade modified trypsin | Promega | Cat. No. V5113 |
| **Critical commercial assays** |  |  |
| Autokit c, glucose diagnostic reagent | Fujifilm Wako | Cat. No. 99703001 |
| HR Series NEFA-HR(2) | Fujifilm Wako | Cat. No. 99934691 |
| RNeasy mini kit | QIAGEN | Cat. No. 74106 |
| Triglyceride Infinity Kit | Thermo Scientific™ | Cat. No. TR22421 |
| β-Hydroxybutyrate (Ketone Body) Colorimetric Assay Kit | Cayman chemical | Cat. No. 700190 |
| **Software and Algorithms** |  |  |
| Adobe Illustrator 2024 | https://adobe.com/products/illustrator | RRID: SCR_010279 |
| GraphPad Prism 10.2.3 | http://www.graphpad.com/scientific-software/prism/ | RRID: SCR_002798 |
| Illumina NovaSeq 6000 | Illumina | NovaSeq 6000 |
| LabView | National Instrument | LabView 8.6 |
| MATLAB | MathWorks | 2022b |
| Origin | Origin Labs | OriginPro 2023 |
| **Deposited data** |  |  |
| Epigenetics data | The GEO repository |  |
| RNA-seq data | The GEO repository |  |
| **Others** |  |  |
| C18 stage tips | Fisher Scientific | Cat. No. 13-110-019 |
| Ketogenic Diet | BioServ | Cat No. S3666 |
| KetoMojo® GK+ Blood Glucose and β-ketone meter | KetoMojo® | GTIN#(01)00858538007615 |
| KetoMojo® GK+ Glucose & Ketone test strips | KetoMojo® | GTIN#(01)00858538007691 |
| Picolab rodent Diet 20 (chow diet) | PicoLab® | Cat No. 5053 |

## RESOURCE AVAILABILITY

### Lead contact

Further information and requests for resources and reagents should be directed to and will be fulfilled by the Lead Contact Dr. Ghazaleh Ashrafi.

### Materials availability

This study did not generate new or unique reagents or other materials.

### Data and code availability

All data reported in this paper will be shared by the lead contacts upon request.

This paper does not report standalone custom code. MATLAB was used to appropriately organize, process, and analyze data and corresponding routines are available from the lead contacts upon request.

Any additional information required to reanalyze the data reported in this paper is available from the lead contacts upon request.

## EXPERIMENTAL MODEL AND STUDY PARTICIPANT DETAILS

### Animals and diets

All animal procedures were in compliance with the US National Institutes of Health Guide for the Care and Use of Laboratory Animals, and conformed to Washington University Animal Studies Committee guidelines. Wild-type male C57BL/6NJ mice (Jackson laboratory, Strain #005304) of 4 and 12 weeks of age were either fed a chow diet (control; PicoLab® Rodent Diet 20) or a high fat (75%), low carbohydrate (3%) Ketogenic Diet (KD, Bio-serv Cat# S3666) for three to four weeks a*d libitum*. Cage bedding was replaced in both groups by aspen chips to avoid any additional nutritional value, and a chewing bone was added in each cage to prevent tooth overgrowth. Weight was monitored once a week for the duration of the diet.

### Blood collection and serum preparation

Glucose and BHB blood levels were monitored weekly and always at the same time of day for the duration of the diet to avoid circadian variability. One drop of blood (∼20uL) was collected from the lateral saphenous vein using a sterile 20G needle following mouse immobilization. Glucose and BHB levels were then immediately analyzed using the KetoMojo® GK+ Blood Glucose and β-ketone Meter and associated strips. For serum analysis, blood was collected by terminal submandibular bleeding on the day of euthanasia with a 21G x 1 ½ Needle. Mice were then immediately euthanized by CO2 overdose and cervical dislocation, and blood was left to clot for two hours at room temperature, and overnight at 4°C. Blood was then centrifuged for 10 minutes at 4°C at 2000xg and supernatant was collected for metabolite analysis.

### Tissue harvesting

Following euthanasia, animal death was ensured by toe pinching. If no reaction was observed, the hippocampal region of the brain was harvested on ice.

### Histone preparation and mass spectrometry

Histone extraction and sample preparation for mass spectrometry were performed as previously described^61^. Brain tissue was dounce homogenized in nuclear isolation buffer (15 mM Tris-HCl, 15 mM sodium chloride, 60 mM potassium chloride, 5 mM magnesium chloride, 1 mM calcium chloride, and 250 mM sucrose at pH 7.5; with freshly added 0.5 mM 4-(2-aminoethyl)benzenesulfonyl fluoride hydrochloride (AEBSF), 10 mM sodium butyrate, 5 mM microcystin, and 1 mM dithiothreitol) with 0.3% CA-630 (Sigma-Aldrich, Cat. No. I8896) and incubated for 5 min on ice. Nuclei were collected by centrifugation (600 × *g* for 5 min at 4°C). The resulting nuclear pellet was washed twice with equal volume of nuclear isolation buffer (without CA-630). Histones were then acid-extracted with 0.2 N sulfuric acid for 4 hr at 4°C with rotation. The insoluble nuclear debris was pelleted (3,400 × *g* for 5 min at 4°C), and the supernatant was retained. Histone proteins were then precipitated by adding 100% trichloroacetic acid in a 1:4 ratio (v/v) and incubating overnight at 4°C. The pellet was washed with acetone to remove residual acid, and histone proteins were resuspended in 30 µl of 50 nM ammonium bicarbonate (pH 8.0). To derivatize primary and secondary amines, the histone sample was mixed with 15 µl derivatization mix (propionic anhydride and acetonitrile in a 1:3 ratio (v/v)), immediately followed by addition of 7.5 µl ammonium hydroxide to maintain pH 8.0. The sample was incubated for 15 min at room temperature, and the derivatization procedure was repeated once. Samples were then dried and resuspended in 50 mM ammonium bicarbonate and incubated with sequencing-grade modified trypsin (Promega, Cat. No. V5113) in a 1:20 enzyme:sample ratio overnight at room temperature. Following trypsin digestion, derivatization was performed twice to derivatize the N-termini of the peptides. Samples were desalted with C18 stage tips (made using Fisher Scientific, Cat. No. 13-110-019). An LC-MS/MS system consisting of a Vanquish Neo UHPLC coupled to an Orbitrap Q Exactive or Ascend (Thermo Scientific) was used for peptide analysis. Histone peptide samples were maintained at 7°C on a sample tray in LC. Separation of peptides was carried out on an Easy-Spray™ PepMap™ Neo nano-column (2 µm, C18, 75 µm x 150 mm) at room temperature with a mobile phase. The chromatography conditions consisted of a linear gradient from 2–32% solvent B (0.1% formic acid in 100% acetonitrile) in solvent A (0.1% formic acid in water) over 48 min and then 42–98% solvent B over 12 min at a flow rate of 300 nL/min. The mass spectrometer was programmed for data-independent acquisition (DIA). One acquisition cycle consisted of a full MS scan and 35 DIA MS/MS scans of 24 m/z isolation width starting from 295 m/z to 1100 m/z. Full MS scans were typically acquired in the Orbitrap mass analyzer across 290–1200 m/z at a resolution of 70,000 or 120,000 in positive profile mode with an injection time of 50 ms and an automatic gain control (AGC) target of 1.0E-06 or 200%. MS/MS data from higher energy collisional dissociation (HCD) fragmentation was collected in the Orbitrap. These scans typically used a nominal collision energy (NCE) of 30 or 25, AGC target of 1000%, and maximum injection time of 60 ms. EpiProfile 3^62^ was used to analyze histone MS data and calculate the ratio at which each modification was present for each peptide (relative abundance).

### Electrophysiology

C57bl6/NJ male mice (4-week-old) were fed a chow diet or a KD for 3 weeks. Hippocampal slices were then prepared as previously described^63^. Briefly, following deep CO_2_ anesthesia, mice were decapitated and brains were dissected out in ice-cold saline containing the following (in mM): 130 NaCl, 24 NaHCO_3_, 3.5 KCl, 1.25 NaH_2_PO_4_, 0.5 CaCl_2_, 5.0 MgCl_2_, and 10 glucose (replacing 5 mM glucose with equimolar BHB in the keto diet animals), pH 7.4 (saturated with 95% O_2_ and 5% CO_2_). Horizontal hippocampal slices (350 μm) were cut using a vibrating microtome (Leica VT1100S)^63^. Slices were initially incubated in the above solution at 35°C for 1 hour for recovery and then kept at room temperature (∼23°C) until use.

Whole-cell patch-clamp recordings using a MultiClamp 700B amplifier (Molecular Devices) in voltage-clamp or current-clamp mode were made from CA1 pyramidal neurons visually identified with infrared video microscopy (Olympus BX50WI; Dage-MTI) and differential interference contrast optics. All the recordings were conducted at near-physiological temperature (33–34°C). The recording electrodes were filled with the following (in mM): 130 K-gluconate, 0.1 EGTA, 2 MgCl_2_, 5 NaCl, 2 ATP-Na_2_, 0.4 GTP-Na, and 10 HEPES, pH 7.3. The extracellular solution contained the following (in mM): 130 NaCl, 24 NaHCO_3_, 3.5 KCl, 1.25 NaH_2_PO_4_, 2 CaCl_2_, 1 MgCl_2_, and 10 glucose (replacing 5 mM glucose with equimolar BHB in the keto diet animals), pH 7.4 (saturated with 95% O2 and 5% CO2). In all experiments, NMDA receptors were blocked with AP-5 (50 μm) to prevent long-term effects. EPSCs (held at −65 mV) and EPSPs were recorded from CA1 pyramidal neurons by stimulating Schaffer collaterals with a monopolar electrode. Synaptic gain was recorded in the presence of 5 µM Gabazine to block inhibition. All data were filtered at 2 kHz, digitized at 20 kHz, acquired using custom software written in LabView or Clampex 11.3, and analyzed using programs written in MATLAB.

### Imaging hippocampal slices with scanning electron microscopy

For sample analysis via Large-Area Scanning Electron Microscopy (LaSEM), mouse brain samples were fixed in a solution containing 2.5% glutaraldehyde and 2% paraformaldehyde in 0.15 M cacodylate buffer with 2 mM CaCl2 pH 7.4. 100 μm coronal vibratome sections of the brains were then stained according to the methods described by Deerinck et. al^64^. In brief, sections were rinsed in cacodylate buffer 3 times for 10 minutes each and subjected to a secondary fixation for one hour in 2% osmium tetroxide/1.5% potassium ferrocyanide in cacodylate buffer for one hour, rinsed in ultrapure water 3 times for 10 minutes each, and stained in an aqueous solution of 1% thiocarbohydrazide for one hour. After this, the tissues were once again stained in aqueous 2% osmium tetroxide for one hour, rinsed in ultrapure water 3 times for 10 minutes each, and stained overnight in 1% uranyl acetate at 4°C. The samples were then washed in ultrapure water 3 times for 10 minutes each and en bloc stained for 30 minutes with 20 mM lead aspartate at 60°C. After staining was complete, coverslips were briefly washed in ultrapure water (3×5 min), dehydrated in a graded acetone series (50%, 70%, 90%, 100% x2) for 10 minutes in each step, and infiltrated with microwave assistance (Pelco BioWave Pro, Redding, CA) into Durcupan resin. Samples were flat embedded between Teflon coated slides and cured in an oven at 60°C for 48 hours.

Post resin curing, sample regions of interest (hippocampus) were selected, removed from the slide with a razor blade, and mounted on a blank block with cyanoacrylate. Thin sections 90 nm in thickness were cut and placed onto 10 mm square silicon wafer chips (Ted Pella, Redding, CA). These chips were then adhered to SEM pins with silver paint and large areas (∼ 330 x 330 µm) were then imaged at high resolution in a FE-SEM (Zeiss Merlin, Oberkochen, Germany) using the ATLAS (Fibics, Ottowa, Canada) scan engine to tile large regions of interest. High-resolution tiles were captured at 16,384 x 16,384 pixels at 5 nm/pixel with a 5 µs dwell time and line average of 2. The SEM was operated at 8 KeV and 900 pA using the solid-state backscatter detector. Tiles were aligned and exported using ATLAS 5.

## QUANTIFICATION AND STATISTICAL ANALYSIS

### Analysis of CA3-CA1 synapses in electron micrographs

Individual synapses were selected from stratum radiatum of the CA1 region of the hippocampus, as neurons in this layer are known to receive input from the Schaffer collaterals from the CA3 region. Synapses with clearly visible active zone (AZ) and post-synaptic density (PSD) were selected from slices prepared from three chow-diet (control) and three KD-fed mice for randomization and further analysis by experimenters blinded to sample type. Active zone length, number of “docked vesicles” within 50nm of the active zone, and total number of synaptic vesicles were determined using Image J.

### Serum metabolite analysis

Total non-esterified fatty acid (NEFA), triglycerides (TG), and glucose levels were analyzed in the mouse serum by the Washington University NORC, Nutrition Obesity Research Center. Serum β-hydroxybutyrate levels were measured using the β-hydroxybutyrate colorimetric assay (Cayman chemicals, #700190) according to the manufacturer’s instructions. Briefly, sera and standards were incubated in the dark for 30min at room temperature with the corresponding reagent. Sample concentrations were determined from the corresponding standard curve and corrected for blanks.

### Electrophysiology

EPCSs during the stimulus trains were normalized to an average of five low-frequency (0.2 Hz) control stimuli preceding each train, to give relative changes in synaptic strength. Each stimulus train was presented four to six times in each cell, and each presentation was separated by ∼2 min of low-frequency (0.2–0.1 Hz) control stimuli to allow complete EPSC recovery to the baseline. To correct for the overlap of EPSCs at short interspike intervals (ISIs), a normalized template of EPSC waveform was created by averaging EPSCs from the same cell with ISI >100 ms. Every EPSC in the train then was approximated by a template waveform scaled to the peak of the current EPSC, and its contribution to synaptic response was digitally subtracted. For determination of paired-pulse ratio (PPR), two EPSCs were recorded at an inter-pulse interval of 25 ms or 10 ms. PPR is the ratio of EPSC2 to EPSC1 amplitude. For measurement of EPSP temporal summation, EPSPs were recorded in response to a train of 5 stimuli at 20, 40 or 100 Hz. EPSP summation is expressed as the percentage increase in EPSP amplitude of 2^nd^ to 5^th^ EPSP relative to the 1^st^ EPSP, i.e., calculated as 100×[(EPSPn – EPSP1)/EPSP1], where EPSP1 is the amplitude of the 1^st^ EPSP and EPSPn is the amplitude of the n-^th^ EPSP in the train.

### RNA sequencing and analysis

Hippocampi were harvested as previously described. Following RNA extraction, samples were prepared according to library kit manufacturer’s protocol, indexed, pooled, and sequenced on an Illumina NovaSeq 6000. Basecalls and demultiplexing were performed with Illumina’s bcl2fastq software and a custom python demultiplexing program with a maximum of one mismatch in the indexing read. RNA-seq reads were then aligned to the Ensembl release 101 primary assembly with STAR version 2.7.9a^65^. Gene counts were derived from the number of uniquely aligned unambiguous reads by Subread:featureCount version 2.0.3^66^. Isoform expression of known Ensembl transcripts were quantified with Salmon version 1.5.2^67^. Sequencing performance was assessed for the total number of aligned reads, total number of uniquely aligned reads, and features detected. The ribosomal fraction, known junction saturation, and read distribution over known gene models were quantified with RSeQC version 4.0^68^.

All gene counts were then imported into the R/Bioconductor package EdgeR^69^ and TMM normalization size factors were calculated to adjust for samples for differences in library size. Ribosomal genes and genes not expressed in the smallest group size minus one sample greater than one count-per-million were excluded from further analysis. The TMM size factors and the matrix of counts were then imported into the R/Bioconductor package Limma^70^. Weighted likelihoods based on the observed mean-variance relationship of every gene and sample were then calculated for all samples with the voomWithQualityWeights^71^ function and were fitted using a Limma generalized linear model with additional unknown latent effects as determined by surrogate variable analysis (SVA)^72^. The performance of all genes was assessed with plots of the residual standard deviation of every gene to their average log-count with a robustly fitted trend line of the residuals. Differential expression analysis was then performed to analyze for differences between conditions and the results were filtered for only those genes with Benjamini-Hochberg false-discovery rate adjusted p-values less than or equal to 0.05.

For each contrast extracted with Limma, global perturbations in known Gene Ontology (GO) terms, MSigDb, and KEGG pathways were detected using the R/Bioconductor package GAGE^73^ to test for changes in expression of the reported log 2 fold-changes reported by Limma in each term versus the background log 2 fold-changes of all genes found outside the respective term. The R/Bioconductor package heatmap3^74^ was used to display heatmaps across groups of samples for each GO or MSigDb term with a Benjamini-Hochberg false-discovery rate adjusted p-value less than or equal to 0.05. Perturbed KEGG pathways where the observed log 2 fold-changes of genes within the term were significantly perturbed in a single-direction versus background or in any direction compared to other genes within a given term with p-values less than or equal to 0.05 were rendered as annotated KEGG graphs with the R/Bioconductor package Pathview^75^.

To find the most critical genes, the Limma voomWithQualityWeights transformed log 2 counts-per-million expression data was then analyzed via weighted gene correlation network analysis with the R/Bioconductor package WGCNA^76^. Briefly, all genes were correlated across each other by Pearson correlations and clustered by expression similarity into unsigned modules using a power threshold empirically determined from the data. An eigengene was then created for each de novo cluster and its expression profile was then correlated across all coefficients of the model matrix. Because these clusters of genes were created by expression profile rather than known functional similarity, the clustered modules were given the names of random colors where grey is the only module that has any pre-existing definition of containing genes that do not cluster well with others. These de-novo clustered genes were then tested for functional enrichment of known GO terms with hypergeometric tests available in the R/Bioconductor package clusterProfiler^77^. Significant terms with Benjamini-Hochberg adjusted p-values less than 0.05 were then collapsed by similarity into clusterProfiler category network plots to display the most significant terms for each module of hub genes in order to interpolate the function of each significant module. The information for all clustered genes for each module were then combined with their respective.

## SUPPLEMENTAL FIGURES

**Figure S1. The effects of ketogenic diet on mouse body weight and serum metabolites. (Related to Fig. 1)**

**(A)** Average body weight of 12-week-old mice during a four-week period on chow (black) or ketogenic diet (KD, green). Not significant: p >0.05; *, 0.05≥ p >0.01; **, 0.01≥ p >0.001. Data are represented as mean ± SEM.

**(B-E)** Levels of glucose **(B)**, BHB **(C)**, free fatty acids **(D)** and triglycerides **(E)** in serum of mice at the end of three weeks of the chow diet (grey) or KD (green). Not significant: p >0.05. Data are represented as mean ± SEM.

N = 5 animals per diet.

**Figure S2. Ketogenic diet alters hippocampal gene expression in young mice. (Related to Fig. 1)**

**(A)** Average body weight of four-week-old male mice during a three-week period on chow (black0 or ketogenic diet (KD, green). Not significant: p >0.05; *, 0.05≥ p >0.01; **, 0.01≥ p >0.001. Data are represented as mean ± SEM.

**(B-E)** Levels of glucose **(B)**, BHB **(C)**, free fatty acids **(D)** and triglycerides **(E)** in serum of mice following three weeks of chow diet (grey) or KD(green). Not significant: p >0.05. Data are represented as mean ± SEM.

N = 6 animals per diet.

**(D)** Pie chart showing the proportion of genes downregulated (60%, 38 genes) or upregulated (40%, 25 genes) in KD compared to chow-diet mice.

**(D)** Heatmap showing differential expression of hippocampal genes in mice on chow or the KD. Log2 fold change (Log2 FC) indicates magnitude of gene expression change in the KD group. Blue, downregulated; red, upregulated.

**(D)** Volcano plot of modulated genes in KD compared to chow diet. Significantly modulated genes are highlighted in green (-Log_10_(adjusted p-value) > -Log_10_(0,05)). Example genes associated with synaptic plasticity and neuronal excitability are labeled.

N = 5 animals per diet. Not significant: p >0.05.

**Figure S3. Baseline EPSC and short-term facilitation is not altered in hippocampal synapses of ketogenic mice. (Related to Fig. 3 and 4)**

**(A, B)** Quantification of baseline EPSC amplitudes recorded in CA1 pyramidal cells before stimulation of Schaffer collaterals with a train of 25 stimuli at 40 Hz **(A)**, or a train of 150 stimuli at 100 Hz to deplete the readily releasable vesicle pool (RRP) **(B)**.

**(C)** Sample traces of paired pulse EPSCs at the inter-pulse interval of 25 milliseconds (ms). Stimulation artifacts were removed for clarity. Traces were scaled to their own 1st EPSC amplitude for visual comparison (vertical bar, 50% of 1st EPSC amplitude).

**(D, E)** Pair pulse ratio at 10ms interval **(D)** and 25ms interval **(E)** recorded in CA1 pyramidal cells in response to Schaffer collateral stimulation with a train of 25 stimuli at 40 Hz.

N = 8 cells/ 3 animals per group. Not significant: p >0.05. Data are represented as mean ± SEM.

## REFERENCES

1. Sultana, B., Panzini, M.-A., Veilleux Carpentier, A., Comtois, J., Rioux, B., Gore, G., Bauer, P.R., Kwon, C.-S., Jetté, N., Josephson, C.B., et al. (2021). Incidence and Prevalence of Drug-Resistant Epilepsy: A Systematic Review and Meta-analysis. Neurology 96, 805–817. 10.1212/WNL.0000000000011839.

2. Rubio, C., López-Landa, A., Romo-Parra, H., and Rubio-Osornio, M. (2025). Impact of the Ketogenic Diet on Neurological Diseases: A Review. Life Basel Switz. 15, 71. 10.3390/life15010071.

3. Tidman, M.M., White, D.R., and White, T.A. (2024). Impact of a keto diet on symptoms of Parkinson’s disease, biomarkers, depression, anxiety and quality of life: a longitudinal study. Neurodegener. Dis. Manag. 14, 97–110. 10.1080/17582024.2024.2352394.

4. Zhang, X., Valeri, J., Eladawi, M.A., Gisabella, B., Garrett, M.R., Vallender, E.J., McCullumsmith, R., Pantazopoulos, H., and O’Donovan, S.M. (2024). Differentially Altered Metabolic Pathways in the Amygdala of Subjects with Schizophrenia, Bipolar Disorder and Major Depressive Disorder. MedRxiv Prepr. Serv. Health Sci., 2024.04.17.24305854. 10.1101/2024.04.17.24305854.

5. Garner, S., Barkus, E., and Kraeuter, A.-K. (2024). Positive and negative schizotypy personality traits are lower in individuals on ketogenic diet in a non-clinical sample. Schizophr. Res. 270, 423–432. 10.1016/j.schres.2024.07.010.

6. Shaafi, S., Mahmoudi, J., Pashapour, A., Farhoudi, M., Sadigh-Eteghad, S., and Akbari, H. (2014). Ketogenic Diet Provides Neuroprotective Effects against Ischemic Stroke Neuronal Damages. Adv. Pharm. Bull. 4, 479–481. 10.5681/apb.2014.071.

7. Guo, M., Wang, X., Zhao, Y., Yang, Q., Ding, H., Dong, Q., Chen, X., and Cui, M. (2018). Ketogenic Diet Improves Brain Ischemic Tolerance and Inhibits NLRP3 Inflammasome Activation by Preventing Drp1-Mediated Mitochondrial Fission and Endoplasmic Reticulum Stress. Front. Mol. Neurosci. 11, 86. 10.3389/fnmol.2018.00086.

8. Paoli, A., Tinsley, G.M., Mattson, M.P., Vivo, I.D., Dhawan, R., and Moro, T. (2024). Common and divergent molecular mechanisms of fasting and ketogenic diets. Trends Endocrinol. Metab. 35, 125–141. 10.1016/j.tem.2023.10.001.

9. Dhatariya, K.K., Glaser, N.S., Codner, E., and Umpierrez, G.E. (2020). Diabetic ketoacidosis. Nat. Rev. Dis. Primer 6, 1–20. 10.1038/s41572-020-0165-1.

10. Nelson, A.B., Queathem, E.D., Puchalska, P., and Crawford, P.A. (2023). Metabolic Messengers: ketone bodies. Nat. Metab. 5, 2062–2074. 10.1038/s42255-023-00935-3.

11. Gilbert, D.L., Pyzik, P.L., and Freeman, J.M. (2000). The ketogenic diet: seizure control correlates better with serum beta-hydroxybutyrate than with urine ketones. J. Child Neurol. 15, 787–790. 10.1177/088307380001501203.

12. Caraballo, R., Vaccarezza, M., Cersósimo, R., Rios, V., Soraru, A., Arroyo, H., Agosta, G., Escobal, N., Demartini, M., Maxit, C., et al. (2011). Long-term follow-up of the ketogenic diet for refractory epilepsy: Multicenter Argentinean experience in 216 pediatric patients. Seizure 20, 640–645. 10.1016/j.seizure.2011.06.009.

13. Ruan, H.-B., and Crawford, P.A. (2018). Ketone bodies as epigenetic modifiers. Curr. Opin. Clin. Nutr. Metab. Care 21, 260–266. 10.1097/MCO.0000000000000475.

14. Düking, T., Spieth, L., Berghoff, S.A., Piepkorn, L., Schmidke, A.M., Mitkovski, M., Kannaiyan, N., Hosang, L., Scholz, P., Shaib, A.H., et al. (2022). Ketogenic diet uncovers differential metabolic plasticity of brain cells. Sci. Adv. 8, eabo7639. 10.1126/sciadv.abo7639.

15. Jensen, N.J., Wodschow, H.Z., Nilsson, M., and Rungby, J. (2020). Effects of Ketone Bodies on Brain Metabolism and Function in Neurodegenerative Diseases. Int. J. Mol. Sci. 21, 8767. 10.3390/ijms21228767.

16. Ma, W., Berg, J., and Yellen, G. (2007). Ketogenic diet metabolites reduce firing in central neurons by opening K(ATP) channels. J. Neurosci. Off. J. Soc. Neurosci. 27, 3618– 3625. 10.1523/JNEUROSCI.0132-07.2007.

17. Juge, N., Gray, J.A., Omote, H., Miyaji, T., Inoue, T., Hara, C., Uneyama, H., Edwards, R.H., Nicoll, R.A., and Moriyama, Y. (2010). Metabolic Control of Vesicular Glutamate Transport and Release. Neuron 68, 99–112. 10.1016/j.neuron.2010.09.002.

18. Hu, E., Du, H., Zhu, X., Wang, L., Shang, S., Wu, X., Lu, H., and Lu, X. (2018). Beta-hydroxybutyrate Promotes the Expression of BDNF in Hippocampal Neurons under Adequate Glucose Supply. Neuroscience 386, 315–325. 10.1016/j.neuroscience.2018.06.036.

19. Qiao, Y.-N., Li, L., Hu, S.-H., Yang, Y.-X., Ma, Z.-Z., Huang, L., An, Y.-P., Yuan, Y.-Y., Lin, Y., Xu, W., et al. (2024). Ketogenic diet-produced β-hydroxybutyric acid accumulates brain GABA and increases GABA/glutamate ratio to inhibit epilepsy. Cell Discov. 10, 1–20. 10.1038/s41421-023-00636-x.

20. Yudkoff, M., Daikhin, Y., Nissim, I., Lazarow, A., and Nissim, I. (2004). Ketogenic diet, brain glutamate metabolism and seizure control. Prostaglandins Leukot. Essent. Fatty Acids 70, 277–285. 10.1016/j.plefa.2003.07.005.

21. Koppel, S.J., Pei, D., Wilkins, H.M., Weidling, I.W., Wang, X., Menta, B.W., Perez-Ortiz, J., Kalani, A., Manley, S., Novikova, L., et al. (2021). A ketogenic diet differentially affects neuron and astrocyte transcription. J. Neurochem. 157, 1930–1945. 10.1111/jnc.15313.

22. Liśkiewicz, D., Liśkiewicz, A., Nowacka-Chmielewska, M.M., Grabowski, M., Pondel, N., Grabowska, K., Student, S., Barski, J.J., and Małecki, A. (2021). Differential Response of Hippocampal and Cerebrocortical Autophagy and Ketone Body Metabolism to the Ketogenic Diet. Front. Cell. Neurosci. 15, 733607. 10.3389/fncel.2021.733607.

23. Li, C., Ma, Y., Chai, X., Feng, X., Feng, W., Zhao, Y., Cui, C., Wang, J., Zhao, S., and Zhu, X. (2024). Ketogenic diet attenuates cognitive dysfunctions induced by hypoglycemia via inhibiting endoplasmic reticulum stress-dependent pathways. Food Funct. 15, 1294– 1309. 10.1039/D3FO04007K.

24. Schwartzkroin, P.A. (1994). Role of the hippocampus in epilepsy. Hippocampus 4, 239–242. 10.1002/hipo.450040302.

25. Malmgren, K., and Thom, M. (2012). Hippocampal sclerosis—Origins and imaging. Epilepsia 53, 19–33. 10.1111/j.1528-1167.2012.03610.x.

26. Kwan, P., Schachter, S.C., and Brodie, M.J. (2011). Drug-Resistant Epilepsy. N. Engl. J. Med. 365, 919–926. 10.1056/NEJMra1004418.

27. Hargreaves, M., and Cameron-Smith, D. (2002). Exercise, diet, and skeletal muscle gene expression. Med. Sci. Sports Exerc. 34, 1505–1508. 10.1097/00005768-200209000-00017.

28. Goldberg, E.L., Shchukina, I., Asher, J.L., Sidorov, S., Artyomov, M.N., and Dixit, V.D. (2020). Ketogenesis activates metabolically protective γδ T cells in visceral adipose tissue. Nat. Metab. 2, 50–61. 10.1038/s42255-019-0160-6.

29. Kennedy, A.R., Pissios, P., Otu, H., Xue, B., Asakura, K., Furukawa, N., Marino, F.E., Liu, F.-F., Kahn, B.B., Libermann, T.A., et al. (2007). A high-fat, ketogenic diet induces a unique metabolic state in mice. Am. J. Physiol.-Endocrinol. Metab. 292, E1724–E1739. 10.1152/ajpendo.00717.2006.

30. Egervari, G., Glastad, K.M., and Berger, S.L. (2020). Food for thought. Science 370, 660–662. 10.1126/science.abb4367.

31. Li, X., Egervari, G., Wang, Y., Berger, S.L., and Lu, Z. (2018). Regulation of chromatin and gene expression by metabolic enzymes and metabolites. Nat. Rev. Mol. Cell Biol. 19, 563–578. 10.1038/s41580-018-0029-7.

32. Periandri, E.M., Dodson, K.M., Vitorino, F.N., Garcia, B.A., Glastad, K.M., and Egervari, G. (2024). Acetate enhances spatial memory in females via sex-and brain region-specific epigenetic and transcriptional remodeling. BioRxiv Prepr. Serv. Biol., 2024.08.26.609729. 10.1101/2024.08.26.609729.

33. Berger, S.L. (2007). The complex language of chromatin regulation during transcription. Nature 447, 407–412. 10.1038/nature05915.

34. Zovkic, I.B., Paulukaitis, B.S., Day, J.J., Etikala, D.M., and Sweatt, J.D. (2014). Histone H2A.Z subunit exchange controls consolidation of recent and remote memory. Nature 515, 582–586. 10.1038/nature13707.

35. Reda, A., Hategan, L.A., McLean, T.A.B., Creighton, S.D., Luo, J.Q., Chen, S.E.S., Hua, S., Winston, S., Reeves, I., Padmanabhan, A., et al. (2024). Role of the histone variant H2A.Z.1 in memory, transcription, and alternative splicing is mediated by lysine modification. Neuropsychopharmacol. Off. Publ. Am. Coll. Neuropsychopharmacol. 49, 1285–1295. 10.1038/s41386-024-01817-2.

36. Guillemette, B., Drogaris, P., Lin, H.-H.S., Armstrong, H., Hiragami-Hamada, K., Imhof, A., Bonneil, E., Thibault, P., Verreault, A., and Festenstein, R.J. (2011). H3 lysine 4 is acetylated at active gene promoters and is regulated by H3 lysine 4 methylation. PLoS Genet. 7, e1001354. 10.1371/journal.pgen.1001354.

37. Deshpande, A.J., Deshpande, A., Sinha, A.U., Chen, L., Chang, J., Cihan, A., Fazio, M., Chen, C.-W., Zhu, N., Koche, R., et al. (2014). AF10 regulates progressive H3K79 methylation and HOX gene expression in diverse AML subtypes. Cancer Cell 26, 896–908. 10.1016/j.ccell.2014.10.009.

38. Cheeseman, K., Jannot, G., Lourenço, N., Villares, M., Berthelet, J., Calegari-Silva, T., Hamroune, J., Letourneur, F., Rodrigues-Lima, F., and Weitzman, J.B. (2021). Dynamic methylation of histone H3K18 in differentiating Theileria parasites. Nat. Commun. 12, 3221. 10.1038/s41467-021-23477-2.

39. Trejo-Arellano, M.S., Mahrez, W., Nakamura, M., Moreno-Romero, J., Nanni, P., Köhler, C., and Hennig, L. (2017). H3K23me1 is an evolutionarily conserved histone modification associated with CG DNA methylation in Arabidopsis. Plant J. Cell Mol. Biol. 90, 293–303. 10.1111/tpj.13489.

40. Jurkowska, R.Z., Qin, S., Kungulovski, G., Tempel, W., Liu, Y., Bashtrykov, P., Stiefelmaier, J., Jurkowski, T.P., Kudithipudi, S., Weirich, S., et al. (2017). H3K14ac is linked to methylation of H3K9 by the triple Tudor domain of SETDB1. Nat. Commun. 8, 2057. 10.1038/s41467-017-02259-9.

41. Abbott, L.F., and Regehr, W.G. (2004). Synaptic computation. Nature 431, 796–803. 10.1038/nature03010.

42. Deng, P.-Y., and Klyachko, V.A. (2011). The diverse functions of short-term plasticity components in synaptic computations. Commun. Integr. Biol. 4, 543–548. 10.4161/cib.15870.

43. Schneggenburger, R., Meyer, A.C., and Neher, E. (1999). Released fraction and total size of a pool of immediately available transmitter quanta at a calyx synapse. Neuron 23, 399–409. 10.1016/s0896-6273(00)80789-8.

44. Schikorski, T., and Stevens, C.F. (2001). Morphological correlates of functionally defined synaptic vesicle populations. Nat. Neurosci. 4, 391–395. 10.1038/86042.

45. Zhai, R.G., and Bellen, H.J. (2004). The architecture of the active zone in the presynaptic nerve terminal. Physiol. Bethesda Md 19, 262–270. 10.1152/physiol.00014.2004.

46. Blaise, J.H., Ruskin, D.N., Koranda, J.L., and Masino, S.A. (2015). Effects of a ketogenic diet on hippocampal plasticity in freely moving juvenile rats. Physiol. Rep. 3, e12411. 10.14814/phy2.12411.

47. Dai, L., Peng, C., Montellier, E., Lu, Z., Chen, Y., Ishii, H., Debernardi, A., Buchou, T., Rousseaux, S., Jin, F., et al. (2014). Lysine 2-hydroxyisobutyrylation is a widely distributed active histone mark. Nat. Chem. Biol. 10, 365–370. 10.1038/nchembio.1497.

48. Dmitrieva-Posocco, O., Wong, A.C., Lundgren, P., Golos, A.M., Descamps, H.C., Dohnalová, L., Cramer, Z., Tian, Y., Yueh, B., Eskiocak, O., et al. (2022). β-Hydroxybutyrate suppresses colorectal cancer. Nature 605, 160–165. 10.1038/s41586-022-04649-6.

49. Shimazu, T., Hirschey, M.D., Newman, J., He, W., Shirakawa, K., Le Moan, N., Grueter, C.A., Lim, H., Saunders, L.R., Stevens, R.D., et al. (2013). Suppression of oxidative stress by β-hydroxybutyrate, an endogenous histone deacetylase inhibitor. Science 339, 211–214. 10.1126/science.1227166.

50. Newman, J.C., and Verdin, E. (2014). β-hydroxybutyrate: Much more than a metabolite. Diabetes Res. Clin. Pract. 106, 173–181. 10.1016/j.diabres.2014.08.009.

51. Chueh, A.C., Tse, J.W.T., Tögel, L., and Mariadason, J.M. (2015). Mechanisms of Histone Deacetylase Inhibitor-Regulated Gene Expression in Cancer Cells. Antioxid. Redox Signal. 23, 66–84. 10.1089/ars.2014.5863.

52. Fioravante, D., and Regehr, W.G. (2011). Short-term forms of presynaptic plasticity. Curr. Opin. Neurobiol. 21, 269–274. 10.1016/j.conb.2011.02.003.

53. Kim, S.M., Ganguli, S., and Frank, L.M. (2012). Spatial Information Outflow from the Hippocampal Circuit: Distributed Spatial Coding and Phase Precession in the Subiculum. J. Neurosci. 32, 11539–11558. 10.1523/JNEUROSCI.5942-11.2012.

54. McClain, K., Tingley, D., Heeger, D.J., and Buzsáki, G. (2019). Position–theta-phase model of hippocampal place cell activity applied to quantification of running speed modulation of firing rate. Proc. Natl. Acad. Sci. 116, 27035–27042. 10.1073/pnas.1912792116.

55. Leutgeb, S., Leutgeb, J.K., Barnes, C.A., Moser, E.I., McNaughton, B.L., and Moser, M.-B. (2005). Independent Codes for Spatial and Episodic Memory in Hippocampal Neuronal Ensembles. Science 309, 619–623. 10.1126/science.1114037.

56. Mongillo, G., Barak, O., and Tsodyks, M. (2008). Synaptic theory of working memory. Science 319, 1543–1546. 10.1126/science.1150769.

57. Deng, P.-Y., Sojka, D., and Klyachko, V.A. (2011). Abnormal presynaptic short-term plasticity and information processing in a mouse model of fragile X syndrome. J. Neurosci. Off. J. Soc. Neurosci. 31, 10971–10982. 10.1523/JNEUROSCI.2021-11.2011.

58. Guang, S., Pang, N., Deng, X., Yang, L., He, F., Wu, L., Chen, C., Yin, F., and Peng, J. (2018). Synaptopathology Involved in Autism Spectrum Disorder. Front. Cell. Neurosci. 12, 470. 10.3389/fncel.2018.00470.

59. Acuña-Catalán, D., Shah, S., Wehrfritz, C., Nomura, M., Acevedo, A., Olmos, C., Quiroz, G., Huerta, H., Bons, J., Ampuero, E., et al. (2024). Ketogenic diet administration later in life improves memory by modifying the synaptic cortical proteome via the PKA signaling pathway in aging mice. Cell Rep. Med. 5, 101593. 10.1016/j.xcrm.2024.101593.

60. Ryvlin, P., Kälviäinen, R., Von Raison, F., Giordano, S., Emir, B., and Chatamra, K. (2010). Pregabalin in partial seizures: a pragmatic 21-week, open-label study (PREPS). Eur. J. Neurol. 17, 726–732. 10.1111/j.1468-1331.2009.02916.x.

61. Sidoli, S., Bhanu, N.V., Karch, K.R., Wang, X., and Garcia, B.A. (2016). Complete Workflow for Analysis of Histone Post-translational Modifications Using Bottom-up Mass Spectrometry: From Histone Extraction to Data Analysis. J. Vis. Exp. JoVE, 54112. 10.3791/54112.

62. Yuan, Z.-F., Lin, S., Molden, R.C., Cao, X.-J., Bhanu, N.V., Wang, X., Sidoli, S., Liu, S., and Garcia, B.A. (2015). EpiProfile Quantifies Histone Peptides With Modifications by Extracting Retention Time and Intensity in High-resolution Mass Spectra. Mol. Cell. Proteomics MCP 14, 1696–1707. 10.1074/mcp.M114.046011.

63. Deng, P.-Y., Kumar, A., Cavalli, V., and Klyachko, V.A. (2022). FMRP regulates GABAA receptor channel activity to control signal integration in hippocampal granule cells. Cell Rep. 39, 110820. 10.1016/j.celrep.2022.110820.

64. Deerinck, T., Bushong, E.A., Thor, A., Ellisman, M.H., Deerinck, T.J., Bushong, E., Ellisman, M., Deerinck, T., and Thor, C.A. NCMIR methods for 3D EM: a new protocol for preparation of biological specimens for serial block face scanning electron microscopy.

65. Dobin, A., Davis, C.A., Schlesinger, F., Drenkow, J., Zaleski, C., Jha, S., Batut, P., Chaisson, M., and Gingeras, T.R. (2013). STAR: ultrafast universal RNA-seq aligner. Bioinforma. Oxf. Engl. 29, 15–21. 10.1093/bioinformatics/bts635.

66. Liao, Y., Smyth, G.K., and Shi, W. (2014). featureCounts: an efficient general purpose program for assigning sequence reads to genomic features. Bioinforma. Oxf. Engl. 30, 923– 930. 10.1093/bioinformatics/btt656.

67. Patro, R., Duggal, G., Love, M.I., Irizarry, R.A., and Kingsford, C. (2017). Salmon provides fast and bias-aware quantification of transcript expression. Nat. Methods 14, 417–419. 10.1038/nmeth.4197.

68. Wang, L., Wang, S., and Li, W. (2012). RSeQC: quality control of RNA-seq experiments. Bioinforma. Oxf. Engl. 28, 2184–2185. 10.1093/bioinformatics/bts356.

69. Robinson, M.D., McCarthy, D.J., and Smyth, G.K. (2010). edgeR: a Bioconductor package for differential expression analysis of digital gene expression data. Bioinforma. Oxf. Engl. 26, 139–140. 10.1093/bioinformatics/btp616.

70. Ritchie, M.E., Phipson, B., Wu, D., Hu, Y., Law, C.W., Shi, W., and Smyth, G.K. (2015). limma powers differential expression analyses for RNA-sequencing and microarray studies. Nucleic Acids Res. 43, e47. 10.1093/nar/gkv007.

71. Liu, R., Holik, A.Z., Su, S., Jansz, N., Chen, K., Leong, H.S., Blewitt, M.E., Asselin-Labat, M.-L., Smyth, G.K., and Ritchie, M.E. (2015). Why weight? Modelling sample and observational level variability improves power in RNA-seq analyses. Nucleic Acids Res. 43, e97. 10.1093/nar/gkv412.

72. Leek, J.T., and Storey, J.D. (2007). Capturing heterogeneity in gene expression studies by surrogate variable analysis. PLoS Genet. 3, 1724–1735. 10.1371/journal.pgen.0030161.

73. Luo, W., Friedman, M.S., Shedden, K., Hankenson, K.D., and Woolf, P.J. (2009). GAGE: generally applicable gene set enrichment for pathway analysis. BMC Bioinformatics 10, 161. 10.1186/1471-2105-10-161.

74. Zhao, S., Guo, Y., Sheng, Q., and Shyr, Y. (2014). Advanced heat map and clustering analysis using heatmap3. BioMed Res. Int. 2014, 986048. 10.1155/2014/986048.

75. Luo, W., and Brouwer, C. (2013). Pathview: an R/Bioconductor package for pathway-based data integration and visualization. Bioinforma. Oxf. Engl. 29, 1830–1831. 10.1093/bioinformatics/btt285.

76. Langfelder, P., and Horvath, S. (2008). WGCNA: an R package for weighted correlation network analysis. BMC Bioinformatics 9, 559. 10.1186/1471-2105-9-559.

77. Yu, G., Wang, L.-G., Han, Y., and He, Q.-Y. (2012). clusterProfiler: an R package for comparing biological themes among gene clusters. Omics J. Integr. Biol. 16, 284–287. 10.1089/omi.2011.0118.

78. Bloom, J., Triantafyllidis, A., Quaglieri, A., Burton Ngov, P., Infusini, G., and Webb, A. (2021). Mass Dynamics 1.0: A Streamlined, Web-Based Environment for Analyzing, Sharing, and Integrating Label-Free Data. J. Proteome Res. 20, 5180–5188. 10.1021/acs.jproteome.1c00683.

